# Assessment of Coupled Phase Oscillators-Based Modeling in Swine Brain Connectome

**DOI:** 10.64898/2026.03.27.713751

**Authors:** Ishfaque Ahmed, Morgan H. LaBalle, Moira F. Taber, Sydney E. Sneed, Erin E. Kaiser, Franklin D. West, Taotao Wu, Qun Zhao

**Affiliations:** Department of Physics and Astronomy, University of Georgia, Athens, Georgia, United States of America; BioImaging Research Center, University of Georgia, Athens, Georgia, United States of America; Institute of Physics, University of Sindh, Jamshoro, Sindh, Pakistan; Regenerative Bioscience Center, University of Georgia, Athens, Georgia, United States of America; Department of Animal and Dairy Science, College of Agricultural and Environmental Sciences, University of Georgia, Athens, Georgia, United States of America; Neuroscience Program, Biomedical and Health Sciences Institute, University of Georgia, Athens, Georgia, United States of America; School of Chemical, Materials and Biomedical Engineering, University of Georgia, Athens, Georgia, United States of America; Institute of Bioinformatics, University of Georgia, Athens, Georgia, United States of America

**Keywords:** Kuramoto model, traumatic brain injury, structural networks, functional networks, functional MRI, swine

## Abstract

Linking structural connectivity (SC) to functional connectivity (FC) through mechanistic models remains challenging in network neuroscience. In this study, empirical data of diffusion magnetic resonance imaging (dMRI) and resting-state functional MRI (rs-fMRI) were used to reconstruct SC and FC of a swine connectome. We evaluated a structurally constrained Kuramoto phase-oscillator framework to reproduce resting-state FC and then assessed the model’s sensitivity to traumatic brain injury (TBI) and its longitudinal progression post-TBI. A joint tuning procedure was implemented to calibrate data-informed natural frequencies and global coupling strength. The tuned Kuramoto model was then used to evolve oscillator phases constrained by the SC, followed by a Balloon–Windkessel hemodynamic model. The optimized model produced significant edge-wise correspondence between averaged simulated FC and the empirical FC (r = 0.61, p < 0.001). Graph-theoretical analysis across network densities (30–50%) showed strong agreement for global efficiency, characteristic path length, and clustering coefficient, while modularity and small-worldness exhibited deviations. Longitudinal analysis of the swine TBI dataset revealed modest reductions in structure–function coupling over time but no significant differences across injury severities. These results demonstrate that optimized Kuramoto models can reproduce key functional network features while preserving inter-subject variability.

## Introduction

The brain processes a variety of information related to different tasks through its anatomical architecture. However, it is challenging in neuroscience to understand how large-scale brain dynamics emerge from this underlying anatomical architecture (Lee et al., 2017; Liégeois et al., 2020). The brain is a complex system composed of interacting dynamic units which are interchangeably termed neuronal ensembles, regions of interest (ROIs), nodes, or oscillators. The collective activity of these units give rise to cognition and behavior (Reeves et al., 2023; Sporns, 2007; Sporns, 2016). These oscillators are connected by white matter wiring known as structural connectivity (SC) that produces functionality-based synchronization, viewed as functional connectivity (FC). The anatomical wiring shows relatively stable connections between brain units while functional synchronization may change based on the task or cognition. Typically, SC is derived from diffusion-weighted magnetic resonance imaging (dMRI) and tract tracing techniques (LaBalle et al., 2026; W. Sun et al., 2024). On the other hand, FC is calculated based on the statistical similarity of temporal neural activity signals from electroencephalography (EEG) (Azhar et al., 2017; Ishfaque Ahmed et al., 2017), functional magnetic resonance imaging (fMRI) (Ahmed et al., 2024; Reeves et al., 2023; Reeves et al., 2025), or magnetoencephalography (MEG). Although the functional dynamics or FC emerges from the brain’s underlying wiring, the direct relationship between them is weak (Zimmermann et al., 2018). Strong functional coupling has been demonstrated by brain regions which are not directly connected by white matter pathways and conversely, inherently connected regions may not always display strong functional synchronizations (Damoiseaux & Greicius, 2009; Honey et al., 2007). Bridging this gap requires mechanistic models that mimic inherent anatomical structure coupling among brain units into emergent functional dynamics.

Graph theory is widely used to measure the topological features of brain networks inferred from neuroimaging data (Ahmed et al., 2025; Pirker-Díaz et al., 2024; Rubinov & Sporns, 2010; Sporns, 2016). Characterization of complex networks using graph theory quantifies several neurobiologically meaningful features that can be easily related to behavioral and cognitive data. These features may facilitate understanding of the global and local organization of brain units and how efficiently they can process information. A large number of studies have shown that brain structural and functional network exhibit modular structure and economical small-world topology (Farahani et al., 2019; Meunier et al., 2009; Pirker-Díaz et al., 2024) which facilitate efficient communication across units with efficient wiring cost (Achard & Bullmore, 2007) and resilient to the random or targeted attacks (Achard et al., 2006; Rayfield et al., 2025). Additionally, graph theoretical framework also has the potential to assess disruptions in connectivity caused by neurological conditions such as traumatic brain injury (TBI). Understanding brain network topology and disruptions caused by TBI is critically important for a personalized, accurate, and timely intervention.

Brain connectivity is established based on distinct brain units/oscillators. Synchronization of these oscillators in the brain can be a key indicator of how well the brain can operate in unison to perform a task. In neuroscience, synchronization phenomenon among brain oscillators plays a crucial role and is studied to understand the physical principle of information processing, routing and transfer, precise task execution, and the perturbation in disease and injury (Palmigiano et al., 2017; Petkoski & Jirsa, 2019; Pirker-Díaz et al., 2024). In a well-functioning brain, there is evidence of a balance between integration (i.e., a state of high global synchronization) and segregation (i.e., a state of local or modular structure; low global synchronization) with flexibility to change according to functional needs (Arenas et al., 2008; Chakravartula et al., 2017; Lee et al., 2017; Palmigiano et al., 2017; Petkoski & Jirsa, 2019; Pirker-Díaz et al., 2024). Neurological conditions including Parkinson’s disease (PD), epilepsy, Alzheimer’s disease (AD), schizophrenia, and autism, have been associated with perturbation in oscillator synchronization (Chakravartula et al., 2017; Kazemi & Jamali, 2022; Pirker-Díaz et al., 2024). Most of these pathologies are associated with abnormal connectivity, failure of brains’ regulatory mechanisms and increased global synchrony. In recent years, the use of the Kuramoto model to investigate synchronization phenomenon in brain oscillators has been increasingly adopted (Lee et al., 2017; Rayfield et al., 2025). In addition, the Kuramoto model simulates the surrogate blood oxygenated level dependent (BOLD) signal that is used for FC estimations. There are some advantages of using the Kuramoto model over other models like neuronal mass, mean field, and conductance models. It is a straightforward, self-regulated and powerful framework to investigate large-scale brain dynamics with output directly comparable to FC and is used to reflect perturbations in functional dynamics from injury or other neurological conditions (Breakspear, 2017; Honey et al., 2007; Pirker-Díaz et al., 2024; Rayfield et al., 2025).

In this study, we applied coupled phase oscillators to model the functional dynamics for the normal pig brain and alterations induced by TBI via underlying structural wiring, which was evaluated by using graph theory and neurodynamic metrics. This study was designed to cover multiple aspects, including assessing the feasibility of the Kuramoto model on pigs, searching for best parameters to simulate empirical FC (eFC), and evaluating whether the Kuramoto model reflects disruptions caused by TBI in simulated FC (sFC). We developed FC and SC using 60 selected cortical ROIs, defined as oscillators for the Kuramoto model, from a previously published pig brain atlas (Saikali et al., 2010). The Kuramoto model evolves phases for the system of oscillators and captures the synchronization of the oscillator population over time, based on the information of underlying structural connections. Further, neural activity simulated by the model was translated into the fMRI BOLD signal using a hemodynamic model. Simulated functional dynamics offers a potentially valuable window to understand the brain’s intrinsic organization and longitudinal progression.

## Methods

### Experimental Data Acquisition and Network Construction

#### TBI induction and data acquisition

A cohort of 44 pigs (8 weeks old, 22 intact males and 22 intact females, Sus scrofa domesticus, Landrace crossbreed) were supplied by the University of Georgia Swine Farm. This cohort of pigs was stratified into three sex-balanced experimental groups. The first group (n=12, referred to as sham) only underwent a craniectomy surgical procedure without TBI, as described by Baker et al., (2019). The other two groups (n=16 each), referred to as mild TBI (mTBI) and severe TBI (sTBI), received induced TBI via controlled cortical impact at a velocity of 4 m/s, a dwell time of 400 ms, and a penetration depth of 3 mm or 9 mm, respectively (Ahmed et al., 2025; Baker et al., 2019; Fagan et al., 2024).

MRI was performed on all pigs at 4 time points referenced against the day of TBI induction, including −1 day, +1 day, +63 days, and +119 days, referring to 1 day pre-TBI, then 1-day, 63-days, and 119-days post-TBI. Anesthesia and MRI protocols were described in previously published work Ahmed et al., (2025). Briefly, pigs were anesthetized and monitored throughout the imaging period to maintain physiological stability and acquired the anatomical T1 weighted (T1w) images using 3D fast spoiled gradient echo (FSPGR) sequence (Ahmed et al., 2025; W. Sun et al., 2024). fMRI images were acquired using the gradient echo-planar imaging (EPI) sequence with imaging parameters TR = 3 s, TE = 30 ms, flip angle = 80°. Each pig underwent a rs-fMRI scan for 305 volumes (15 minutes and 15 seconds) with a field of view = 12.8 × 12.8 × 6.2 cm, and an acquisition matrix of 96 × 96 × 31 in a coronal slice orientation (Ahmed et al., 2024; Reeves et al., 2023). A spin-echo EPI sequence with an isotropic resolution of 2 mm was used to record 3 non-diffusion images followed by 30 diffusion gradient images. Acquisition parameters were TR = 10,000 ms, optimized minimum TE typically below 90 ms, flip angle = 90°, acquisition matrix of 96 × 96, 32 slices, and a field of view of 192 mm. Additionally, reversed phase-encoding directions (anterior-to-posterior and posterior-to-anterior) were also acquired to apply corrections to susceptibility-induced distortions. This study was performed in accordance with the National Institutes of Health (NIH) Guide for the Care and Use of Laboratory Animals. All procedures were reviewed and approved by the University of Georgia Institutional Animal Care and Use Committee (Animal Use Protocol: A2023 07-021-Y3-A12).

#### Data preprocessing and network construction

The rs-fMRI and DTI images from the pig dataset were preprocessed in accordance with our earlier publications (Ahmed et al., 2025; W. Sun et al., 2024) prior to network constructions. Briefly, this preprocessing includes data format conversion, head motion corrections, slice time corrections, skull stripping, and spatial normalization to the template space. In this study, a predefined anatomical atlas was used for parcellation into ROIs (Saikali et al., 2010). To ensure fair comparison across anatomical structures, we excluded ROIs with a volume smaller than 100 mm^3^, resulting in 60 ROIs (see a full list in **Supplementary Table 1**). The anatomical atlas was then registered to rs-fMRI and DTI spaces separately using first volume and B0 image respectively. To construct FC matrices, a customized MATLAB script was used to extract time series data for each ROI and estimate correlation across ROIs via Pearson correlation coefficient (Ahmed et al., 2024; Reeves et al., 2025; Sporns, 2016). The self-correlation was ignored by replacing the diagonal with zeros. The FSL probabilistic tractography tool (PROBTRACKX) was used to construct SC matrices of each pig. To provide symmetry, upper and lower triangles of output matrix were then averaged. Each of the FC and SC matrices were normalized by their absolute maximum value.

### FC simulation via coupled oscillators

#### Kuramoto model for coupled oscillators

Neuronal oscillatory mechanism is the fundamental process of information transfer within and across brain regions (Bonnefond et al., 2017). This oscillatory behavior is associated with task or voluntary processes exhibited across multiple spatial and temporal scales. Cortical brain regions (ROIs) or oscillators can communicate to each other by means of the synchronization of these phase oscillations. As an illustration, 4 oscillators are shown (**Figure 1H**) in a pig brain with their phase oscillations at low frequency fluctuations (Bonnefond et al., 2017). We used the Kuramoto model for coupled phase oscillators to simulate the synchrony of low frequency BOLD signals (Kuramoto, 2005; Okuda & Kuramoto, 1991; Pirker-Díaz et al., 2024; Rayfield et al., 2025). Further, each of the oscillators were coupled to others via inherent anatomical connections (as wires depicted in **Figure 1H**), where thickness of the wires indicates strength of coupling.

**Figure 1.**
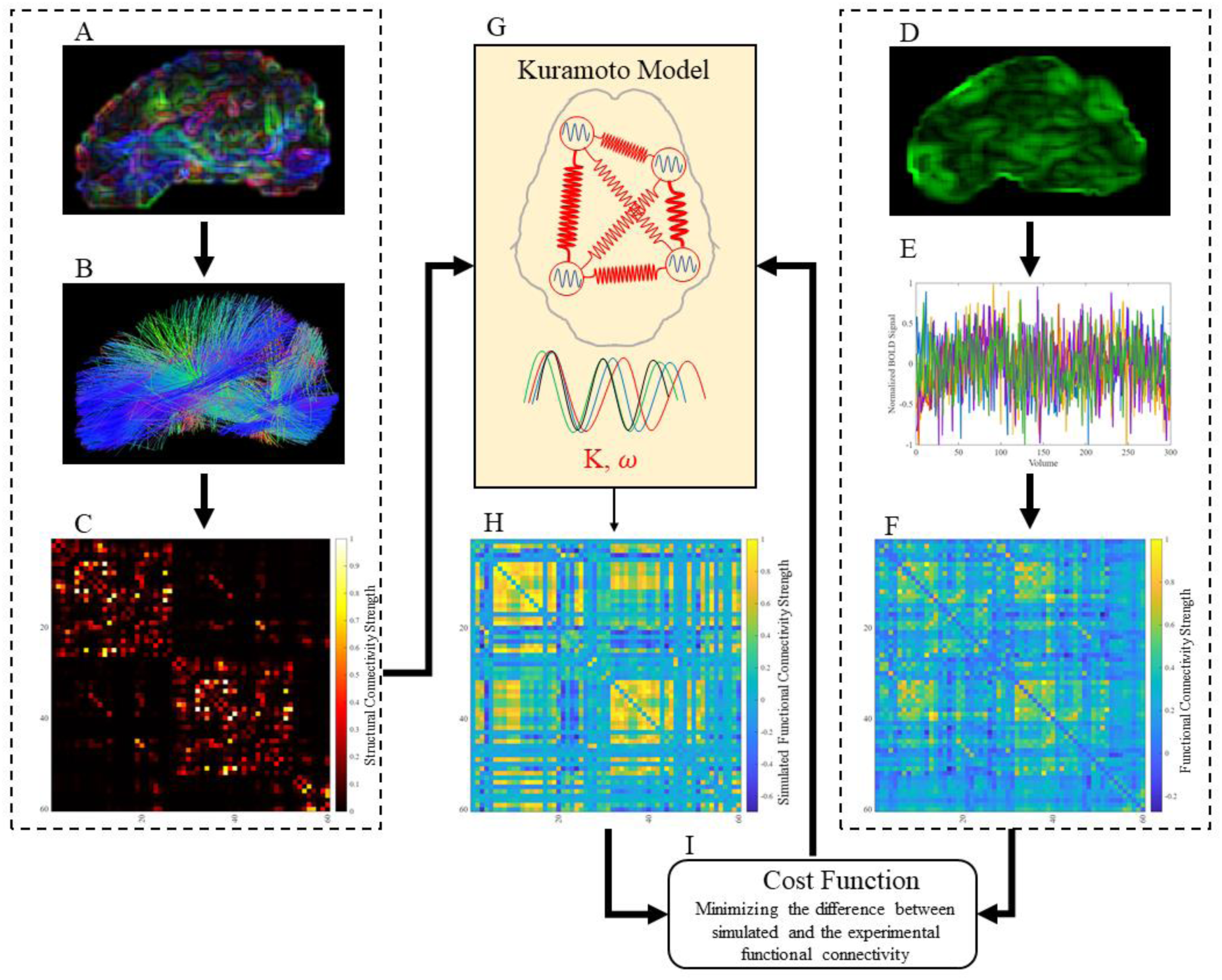
Schematic diagram of the joint optimization framework for global coupling constant and natural frequencies. **Left column**: Panel A illustrates diffusion tensor imaging data; panel B shows a tractogram identified via tractography, panel C manifests the structural connectivity matrix. **Right column**: Panel D shows functional data, panel E shows extracted BOLD signal for each ROI and panel F depicts functional connectivity matrix. **Middle column**: Panel G represents the Kuramoto model framework by illustrating four brain oscillators inherently connected by their local coupling and used to simulate functional patterns shown in panel H. These functional networks are compared by cost function in panel I and coupling constant and natural frequencies are updated in the model presented in panel G.

The phase evolution for each oscillator over time is done by numerically solving a system of equations as described in equation 1 (Achard et al., 2006; Rayfield et al., 2025).

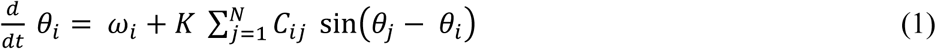

where, *θ*_*i*_ is the list of evolved phases and *ω*_*i*_ is the angular frequency (2*π* times the natural frequency) associated with *ith* oscillator. The *ω* vector may comprise different values which reflect the heterogeneity of the brain regions. *K* represents the global coupling constant and *C*_*ij*_ is the anatomical connection strength between *ith* and all other oscillators as *j* = 1,2,3,…, *N*, as depicted in **Figure 1C**.

Furthermore, inter-regional communication of the brain is not instantaneous. There is always a lag in signal transmission which depends upon the fiber length connecting two ROIs and the conduction speed of the fiber. Incorporation of time delay information makes the Kuramoto model biophysically more plausible and therefore may enhance the correspondence between sFC and eFC (Cabral et al., 2012; Deco et al., 2011). The Kuramoto model incorporating time delays is given by

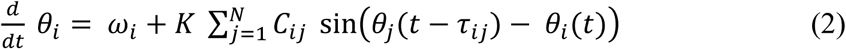

where *τ*_*ij*_ indicates time delay between oscillators *i* and *j*. The delay differential equations were numerically solved in MATLAB (The MathWorks Inc., Natick, MA) following (Wu et al., 2022).

A key feature of the Kuramoto model is the measure of transition of phases from incoherence to partial or fully synchronized state as a function of coupling strength (Cabral et al., 2012; Okuda & Kuramoto, 1991; Strogatz, 2000). This feature is the order parameter *R*(*t*) of the Kuramoto model, given as

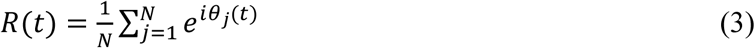

where *R*(*t*) measures the level of synchronization among oscillators and ranges 0 ≤ *R* ≤ 1. A system of oscillators is said to be fully synchronized when *R* = 1 and desynchronized when *R* = 0. The temporal mean of the Kuramoto order parameter quantifies the global synchrony, and its standard deviation measures metastability. Global synchrony provides insights into overall coherence while metastability indicates the flexibility of the system to switch states (Breakspear, 2017; Cabral et al., 2012; Lee et al., 2017). In this study, we used synchrony and metastability as complementary metrics to determine the optimal global coupling constant.

#### Parameter optimization

In a coupled-oscillators framework, natural frequencies of the oscillators and the global coupling constant play critically important roles to simulate the experimental FC. There is no consensus over the natural frequencies range; however, normally distributed frequencies with a mean of 60 Hz and standard deviation of 1 Hz were predominantly adopted (Cabral et al., 2012; Lee et al., 2017; Wu et al., 2022). A few other studies have found frequencies < 30 Hz more appealing because this range covers the multiple frequency bands associated with electroencephalographic signals (Capilla et al., 2022; Lew et al., 2021). Another option especially dealing with the fMRI data is to use frequencies below Nyquist sampling frequency (Rayfield et al., 2025). It is important to note that real functional dynamics of the brain can be accurately reproduced by the Kuramoto model framework only with parameters correctly reflecting the brain, especially the global coupling constant and the natural frequencies. The use of randomly selected or inappropriately optimized parameters may fail to capture the real functional dynamics of the brain or reproduce sub-optimal or misleading FC patterns. Therefore, we developed an algorithm (**Figure 1**) to calibrate the data-informed natural frequencies of the oscillators in contrast to previously used approaches. In this context, a joint optimization criterion was applied to experimental FC and SC data over the range of *K* = 1 − 40 as

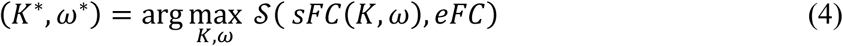

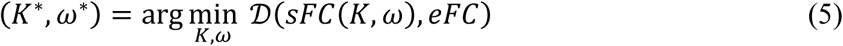

where *eFC* and *sFC* are experimental and simulated FC matrices respectively. *K* represents global coupling constant and *ω* = 2*πf*, where *f* is a vector of natural frequencies. The optimization metrics 𝒮 and 𝒟 are measures of similarity and difference in terms of Pearson correlation and mean squared error. The frequencies were initialized randomly between 0 and 1, then updated iteratively based on the measured difference between sFC and eFC to maximize similarity and minimize difference. A detailed block diagram is shown in **Supplementary Figure S1**, which was used to optimize the parameters over a certain range of coupling constants. The optimal values of *K* and *f* were chosen by a grid selection from joint similarity and difference measures. It was achieved by an iterative procedure of selecting *x*% top grids from similarity matrix and lower *x*% from difference matrix. This process continued until an overlapping grid was identified, where the first overlapping grid represents the optimal value of *K* and the natural frequencies.

#### Simulating FC networks

The Kuramoto model described in Equation 1 was used to simulate FC via dynamic phase coherence in a tuning process of parameters to save computational cost. However, a more realistic model to reproduce FC should incorporate hemodynamic delays (Friston et al., 2000) and translate phases into BOLD signal. The Balloon-Windkessel hemodynamic model (Wu et al., 2022) was incorporated into the framework to transform sine of phase into BOLD signal. Like the empirical fMRI data, we simulated BOLD signals for a time of period (970 seconds) with the same temporal resolution as the experimental data. We discarded the first 70 seconds of BOLD signal as a standard procedural step to remove non-steady-state signal (Ahmed et al., 2024). The sFC was then calculated via simulated BOLD signal using Pearson correlation. Following this pipeline, we predicted sFC for all experimental SC datasets, covering all normal control, sham, and TBI groups across the full set of time points using corresponding empirical SC networks.

#### Null model testing and network-based analysis

Once the global coupling constant and natural frequencies were optimized, we evaluated the Kuramoto oscillators to determine if dynamic patterns and FC network properties were being captured. In this context, a null model testing was performed following (Rayfield et al., 2025). Under this null model, we simulated sFC for all normal control pigs using 2 approaches: 1) empirical SC and 2) randomized empirical SC. The randomized SC networks were generated by shuffling the edges of empirical SC networks while preserving the network density and mean degree (Rubinov & Sporns, 2010). The sFCs were then compared against eFC via Pearson correlation to measure how similar FC networks were reproduced by the model. Finally, recorded similarity score distribution from randomized (null model) SC approach was statistically evaluated against similarity score distribution from empirical SC approach to assess whether the Kuramoto model simulated FC patterns that showed greater similarity to empirical FC than those generated using null models.

### Graph theory analysis

We calculated the graph theoretical metrics for eFC and sFC networks. These networks were first transformed into undirected binary networks by applying thresholding. Instead of extracting graph theoretical features at an arbitrary threshold, we constructed adjacency networks at multiple connection densities ranging from 1–100%. Comparison between topological features of empirical and simulated graphs at same densities throughout the range has demonstrated greater reliability (Achard & Bullmore, 2007; Lee et al., 2017; Van Wijk et al., 2010). Graph topological features were observed to show randomness at very low connection densities due to the fact that graph may comprise multiple fragments (Fallani et al., 2007; Lynall et al., 2010). In this context, Fallani et al., (2007) established that network degree and other graph features stabilize at 30% connection density, while Lynall et al., (2010) determined fully connected networks with stable features at 37–50% connection densities. This motivated us to look further into the comparison of the features ranging from 30–50% connection densities. The graph features investigated to compare eFC and sFC included global efficiency (GE), characteristic path length (CPL), modularity (MOD), small world-ness (SW), and global clustering coefficient (CC) (see Appendix A for definitions). In this study, we complemented the overlay of graph theory metrics with a deeper assessment of the differences through relative error (Lee et al., 2017) analysis between eFCs and sFCs. The relative error was measured using empirical values as the baseline in equation 6

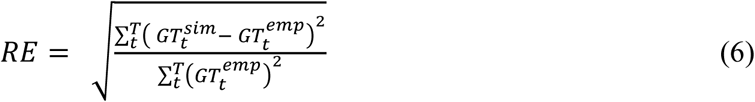

where 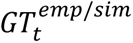 indicate graph theory metric measured from empirical and simulated networks, *t* represents the starting and *T* represents final threshold density value with a unity resolution.

### Longitudinal evaluation of the Kuramoto model

To examine how well the model generalizes across time, we applied the tuned model based on normal control SC networks of pigs to predict FC at the +1 day, +63 days, and +119 days time points using corresponding SC networks of pigs. Longitudinal evaluation included both network-level comparisons between sFC and eFC networks and graph-theoretical metric comparisons. In the network-level evaluation, Pearson correlation coefficients were calculated between the sFCs and eFCs (excluding diagonal elements of the matrices) for each pig. This evaluation provided insights into the following 4 questions: 1) how well the model simulates the functional dynamics and FC patterns; 2) whether it preserves the heterogeneity of the data; 3) whether the TBI injury and its severity affects the model’s ability to reproduce FC; and 4) whether tuned parameters work well for a longitudinal study. On the other hand, graph-theoretical comparison provided insights into how accurately the model reproduces key network features. In longitudinal assessment, we quantified the relative error for graph-theoretical features obtained from sFC and eFC matrices.

## Results

### Optimization of natural frequencies and coupling constant

In this study, we optimized the natural frequencies and global coupling constant using functional and structural matrices of the control group (44 healthy pigs, **Table 1**) pigs individually and the average of all pigs. **Figure 2(A–C)** presented measures of similarity, difference, and optimized natural frequencies based on the optimization procedure of individual pig connectivity matrices. The heatmaps in **2A** and **2B** are the average of 44 individually optimized similarity and difference maps, while the bar plot in **2C** depicts mean and standard deviation of the optimized natural frequencies of all 60 oscillators. As a comparison, **Figure 2(D–F)** showed the same results using an averaged FC and SC matrix of all 44 pigs. The scattered plot in F showed the distribution of natural frequencies. The natural frequencies in both **Figures 2C** and **2F** were color-coded into 3 sections: left hemisphere (ROIs #1-26), right hemisphere (ROIs #27-52), and ROIs associated cerebellum (ROIs #53-60).

**Figure 2.**
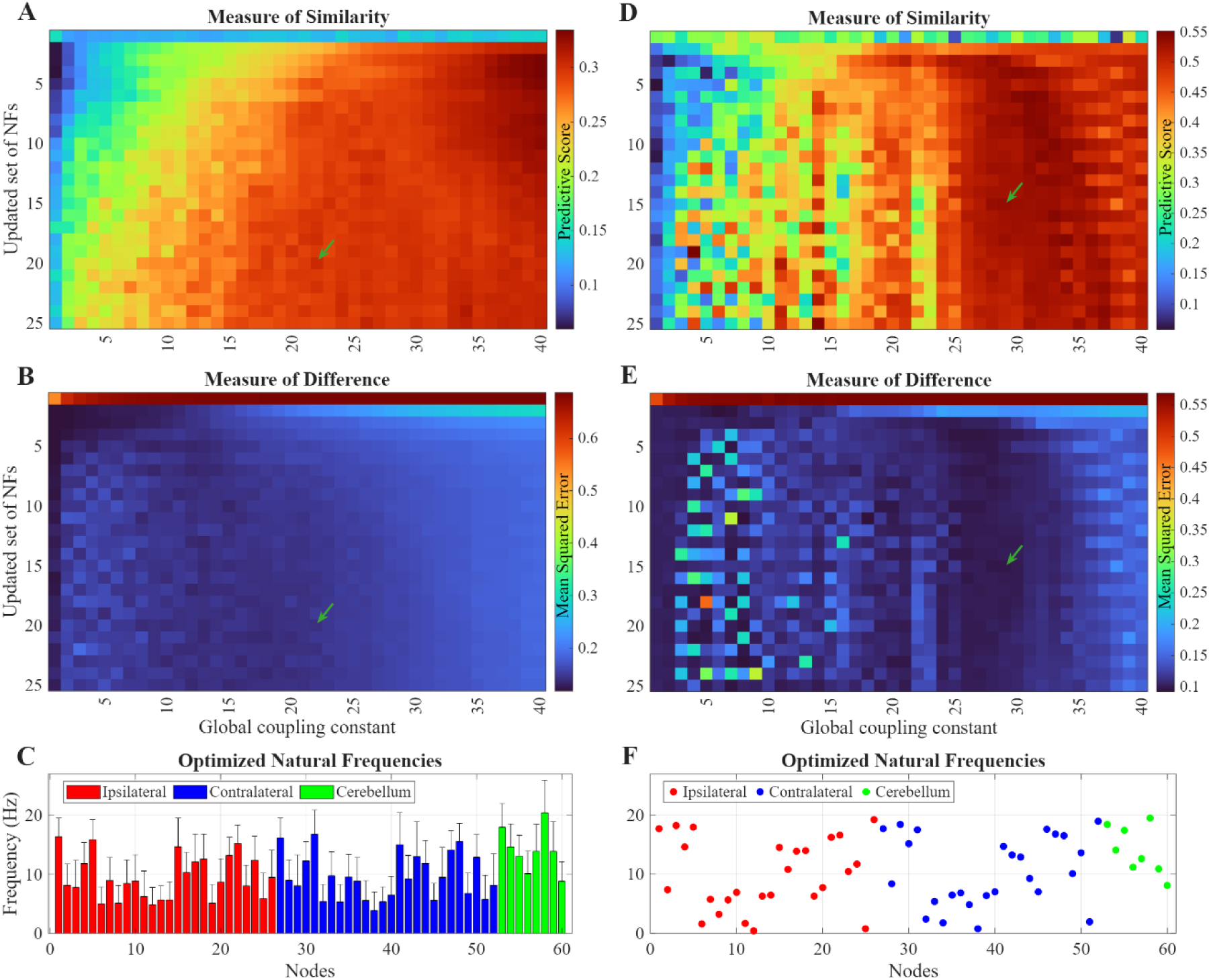
Depicts the measure of similarity, measure of difference and distribution of natural frequencies. **Left column**: results are based on optimization using individual pigs. Panel A shows a heatmap of correlations; heatmap in B shows mean squared error between simulated and empirical functional connectivity. Panel C shows the distribution of natural frequencies for all oscillators. **Right column**: results are based on optimization using averaged matrix of all control pigs. Panel D shows a heatmap of correlations, heatmap in E shows mean squared error between simulated and empirical functional connectivity. Panel F shows the distribution of natural frequencies for all oscillators. The optimal *K*, selected by the individual and averaged procedures with higher correlations and lower mean squared errors are 22 and 29, as indicated by the arrows.

**Table 1.** Number of pigs in the groups at each time point.

| Group | -1 day | +1 day | +63 days | +119 days |
| --- | --- | --- | --- | --- |
| Normal Control | 44 | - | - | - |
| Sham | - | 12 | 12 | 12 |
| mTBI | - | 16 | 16 | 16 |
| sTBI | - | 16 | 15 | 16 |

It can be seen from similarity and difference measures of both procedures that reproduced FC patterns matched better beyond certain *K* values and showed lower mean squared errors. The optimal *K* selected by the individual and averaged procedures are 22 and 29, respectively. In addition, tuned frequencies from the individual and averaged tuning procedure showed great bilateral similarity (i.e., ROIs #1-26 vs. ROIs #27-52, in **Figure 2C** and **2F**).

### Model evaluation via order parameter

Once the model parameters were optimized, we measured the synchrony and the metastability using values of Kuramoto order parameter against a range of *K*. Since synchrony and metastability can help in identify a balanced coherence and flexibility of the system, we used these measures to validate the optimal *K* and natural frequencies. In **Figure 3A**, synchrony and metastability were presented as a function of *K* based on individual *K* and natural frequencies of the normal control group data (44 pigs). The oscillators had small coherence (< 0.3) at the first few *K* values. Then the coherence increased gradually and ranged between 0.3–0.6, showing that the coherence of system is slowly improving with increasing *K*. **Figure 3B** showed metastability against the same *K* range. An optimal *K* may be selected from a region where system of oscillators was substantially synchronized and had a pronounced metastable dynamics (Wang et al., 2025). Based on the synchrony (**Figure 3A**), an optimal value could be selected as *K* > 20, while the metastability map (**Figure 3B**) suggested 10 < *K* < 23. As a result, the order parameter measurements for individual optimization supported the selection of *K* = 22 as identified in joint measure of similarity and difference. On the other hand, in **Figure 3C** and **3D** (average-based *K* and natural frequency optimization), the synchrony and metastability values were relatively higher than the individual optimization scheme. The mean synchrony curve (**Figure 3C**) flattened at *K* ≥ 20, indicating plateau with an increased stability in synchrony measure, which suggests the optimal values of *K* > 25; while the metastability (**Figure 3D**) favored 15 < *K* < 30. Following the criterion of high synchrony and metastability, a value of *K* = 29 was selected, which is also consistent with findings in the previous sub-section. Additionally, averaged eFC data was found to be cleaner and stable in contrast to individual eFC graphs as shown in **Supplementary Figure S5**. Functional graphs were very heterogenous with similarity value against mean FC is 0.61 ± 0.11 while SC graph were very consistent with similarity against averaged SC graph of 0.97 ± 0.01 (see **Supplementary Figure S6**).

**Figure 3.**
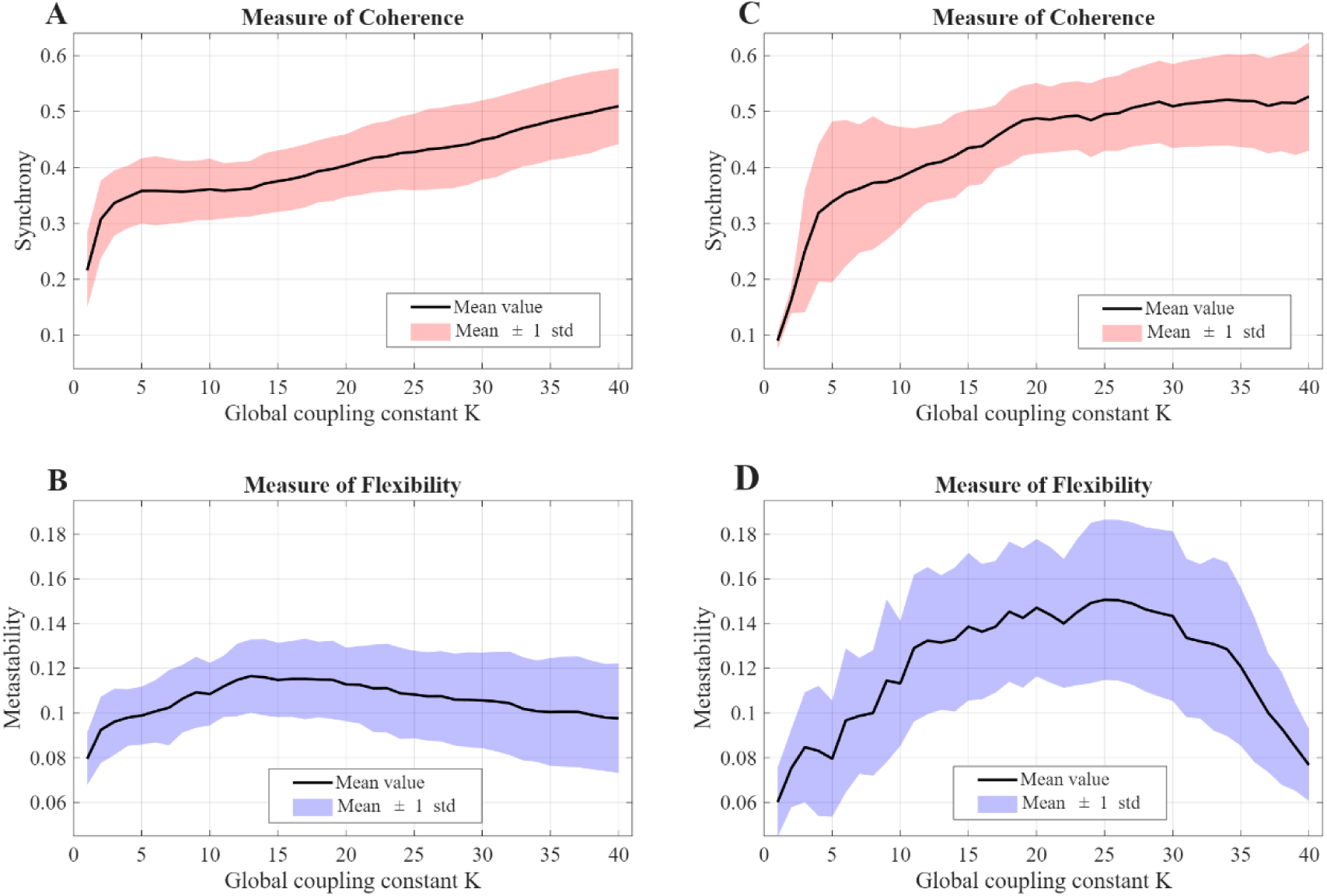
Representation of synchrony and metastability. **Left**: Panels A and B depict synchrony and metastability (mean value and its spread) using individual *K* and natural frequencies. **Right**: Panels C and D depict synchrony and metastability (mean value and its spread) using average-based *K* and natural frequencies.

### Simulated FC Networks

The tuning process yielded 3 possible choices of *K* and natural frequencies including 1) an averaged scheme (Avg) using *K* = 29 and optimized natural frequencies in **Figure 2F**; 2) individual scheme (Indv) using *K* and frequencies from individual pigs (maps as shown in **Supplementary Figure S2** for 2 pig examples); 3) combined Avg/Indv scheme (Combn) using *K* = 29 (from averaged tuning) and pig-specific natural frequencies from individual maps. We then simulated sFC matrices for all control pigs and measured correlation between sFC and eFC matrices for the 3 schemes (**Figure 4A**). The 3 schemes yielded similar distributions of correlation coefficients, with the Combn scheme showing a higher mean than the other two. Adopting individual natural frequencies can potentially preserve the heterogeneity of individual pigs. The Combn scheme with pig-specific natural frequencies and global coupling constant *K* = 29 was therefore used for all following simulations of sFC. Further, a null model test was conducted on selected scheme to verify that sFCs were a manifestation of empirical structure-function relationships, rather than any representation by chance. In this context, sFC matrices were reproduced using original (empirical) and randomized SC (shown in **Supplementary Figure S3**) and were compared against respective eFC matrices. In **Figure 4B**, the similarity score (correlation coefficient) profiles from both original and randomized SC were subjected to a statistical comparison.

**Figure 4.**
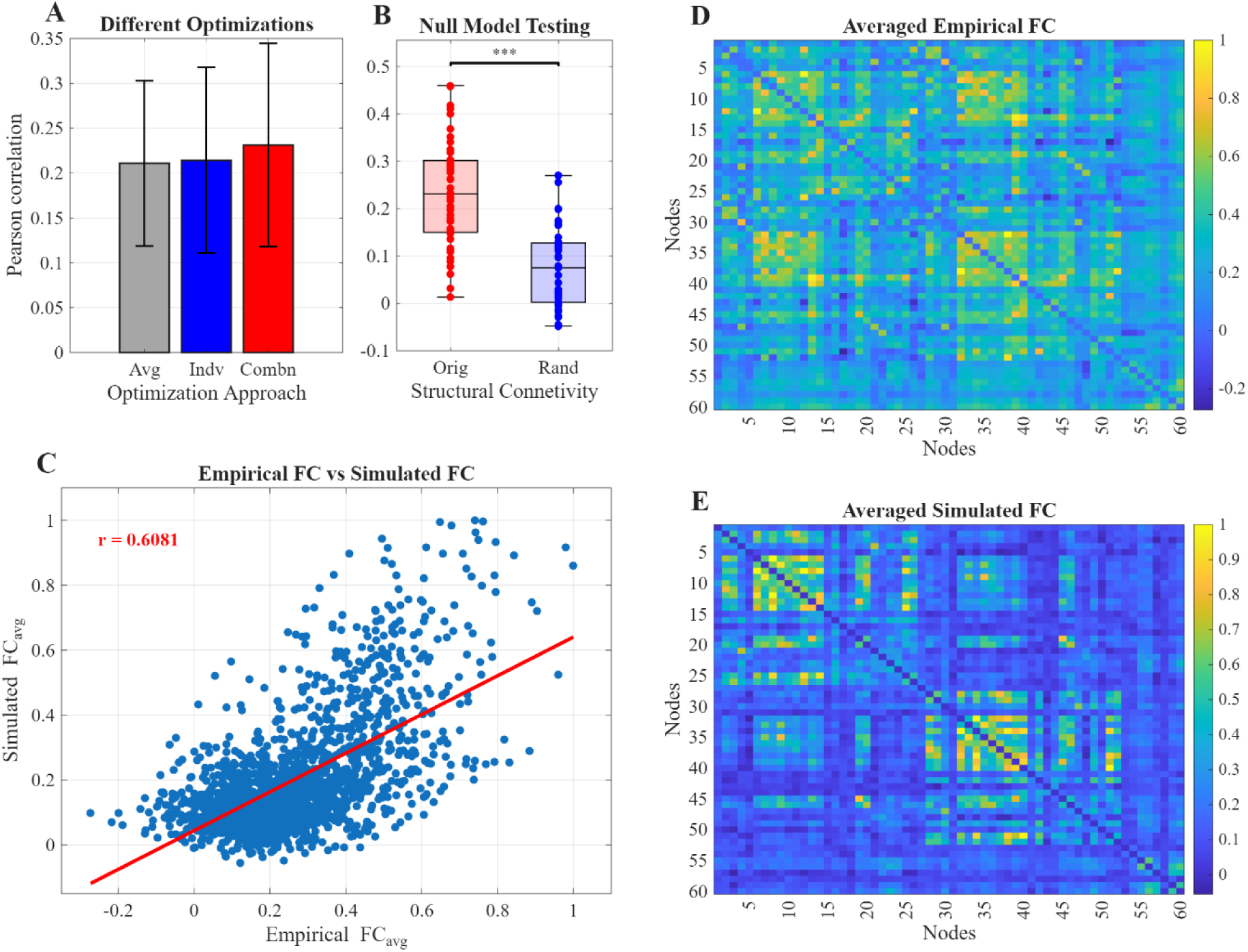
Comparison of the Kuramoto model performances using three optimization schemes and network comparison between simulated FC and empirical FC. Panel A shows the comparison of three approaches, averaged (Avg), individualized (Indv), and combined Avg/Indv (Combn). Panel B indicates the edge-based relationship between averaged simulated and empirical FC. Panel C shows averaged empirical FC graph of all control pigs and panel D shows averaged simulated FC graph.

It is evident from the box charts that sFC using original SC showed significantly higher similarity with eFC. This testing offered 2 key benefits: underlying brain structural organization shapes FC patterns and the Kuramoto model accurately reflected these FC patterns instead of generating arbitrary/random FC patterns. Next, we evaluated the edges (or edge weights) of the averaged eFC 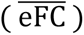 and averaged sFC 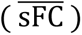. **Figure 4C** presented a scatter plot between the off-diagonal elements/edges of 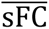 and 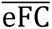. The correlation between the vectors of the edges was found to be r = 0.61 (p < 0.001), showing a strong match between the 2 FC graphs. Further, heatmaps in **Figure 4D** for 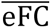 and **4E** for 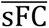 also showed a good resemblance of overall network organizations.

### Graph theory analysis

To further evaluate how closely Kuramoto model reproduces topological features in the simulated networks, we examined 5 global graph theoretical metrics across eFCs and sFCs using the normal control group data (**Table 1**). A stable density range for graph theory metrics should be between 30–50%. **Figure 5(A–E)** showed global graph theory metrics, including GE, CPL, MOD, CC, and SW, within a density range of 30–50% for the normal control (i.e., −1 day) pigs. Comparison of these metrics for an entire range of network densities (1–100%) is shown in **Supplementary Figure S4**. In **Figure 5A**, we observed a close match of GE between eFC and sFC. A relatively good overlap of overall distributions for CPL and CC is presented in **Figure 5B** and **5D**. It is also observed in **Figure 5C** and **5E** that simulated MOD and SW distributions match poorly with their respective empirical distributions. The bar plot presented in **Figure 5F** provides relative errors (average and standard deviation) between simulated and empirical graph theory metrics measured using equation 6. The relative error analysis supported the findings from the overlay of the graph metric distributions in **Figures 5(A–E)** that GE, CPL, and CC features are reproduced relatively better in sFC than MOD and SW.

**Figure 5.**
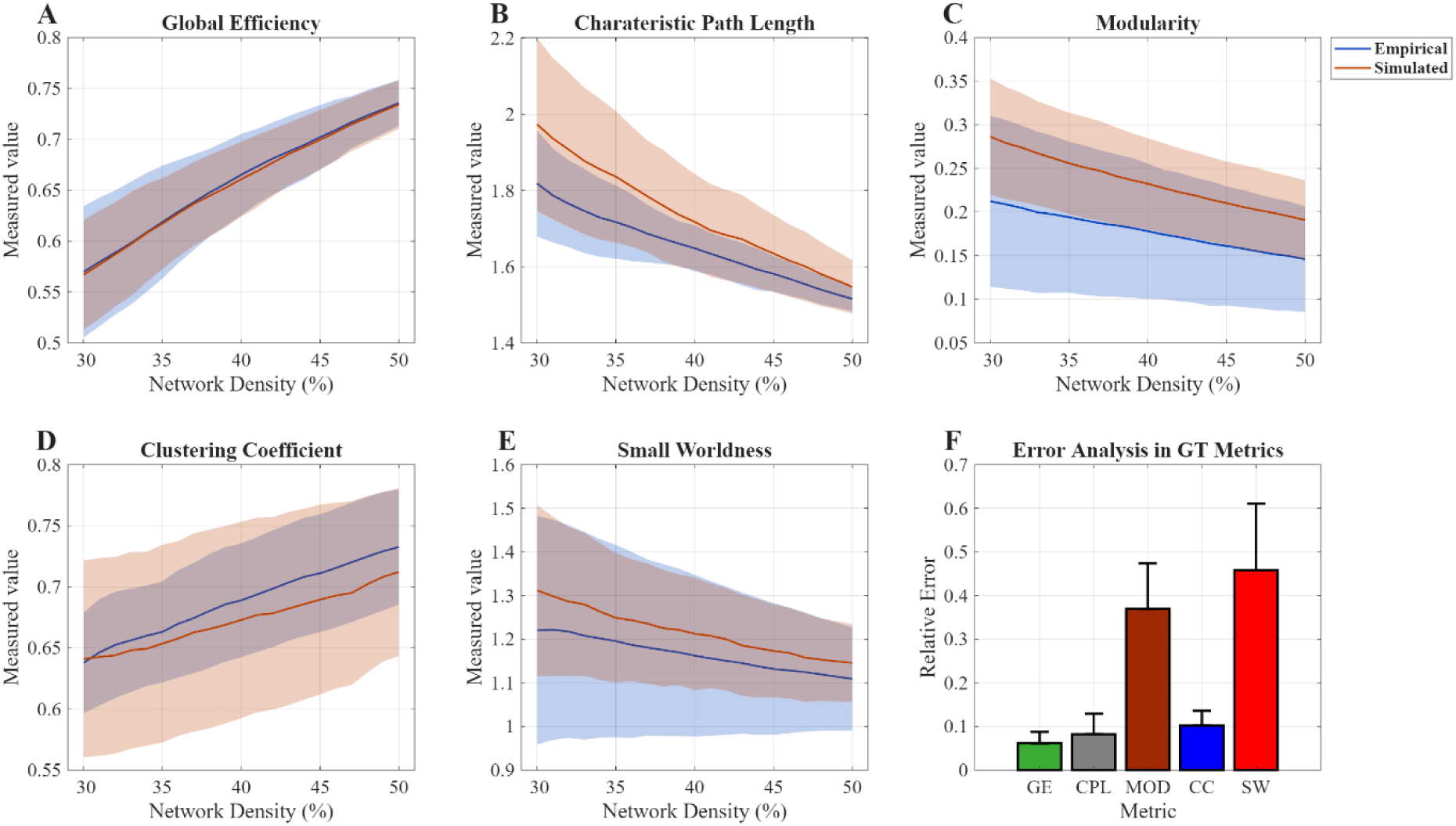
Evaluation of the Kuramoto model based on graph theoretical features in the range of 30–50% network densities. Overlay of the mean and distributions of empirical and simulated global efficiency (panel A), characteristic path length (panel B), modularity (panel C), clustering coefficient (panel D) and small worldness (panel E). Panel F shows the measure of relative error for all the 5 metrics.

### Evaluation of the Kuramoto model using longitudinal TBI data

Extending the optimized Kuramoto model using the normal control data to evaluate our model’s performance for later time points yielded valuable insights, as a good correlation between sFC with eFC revealed brain recovery processes. **Figure 6A** provided an assessment of the model’s ability to predict FC patterns at the later 3 time points. The prediction score (i.e., correlation between simulated and empirical networks) in **Figure 6A** showed a wide-ranged distribution, 0.02–0.47, for the normal control (−1 day, the gray box chart) pigs. Continuing to the +1 day time point, all 3 groups showed slightly lower median values compared to normal control, however the spread of prediction score distributions remained invariant. No substantial differences across groups were identified at +1 day. At +63 days, the prediction score decreased significantly irrespective of injury severity, yet no significant difference was identified across groups. Similarly, prediction score distributions for each group at +119 days seem unchanged with slight variations in median values. This may be an indicator of the stability of inherent structure and properties between +63 days and +119 days time points.

**Figure 6.**
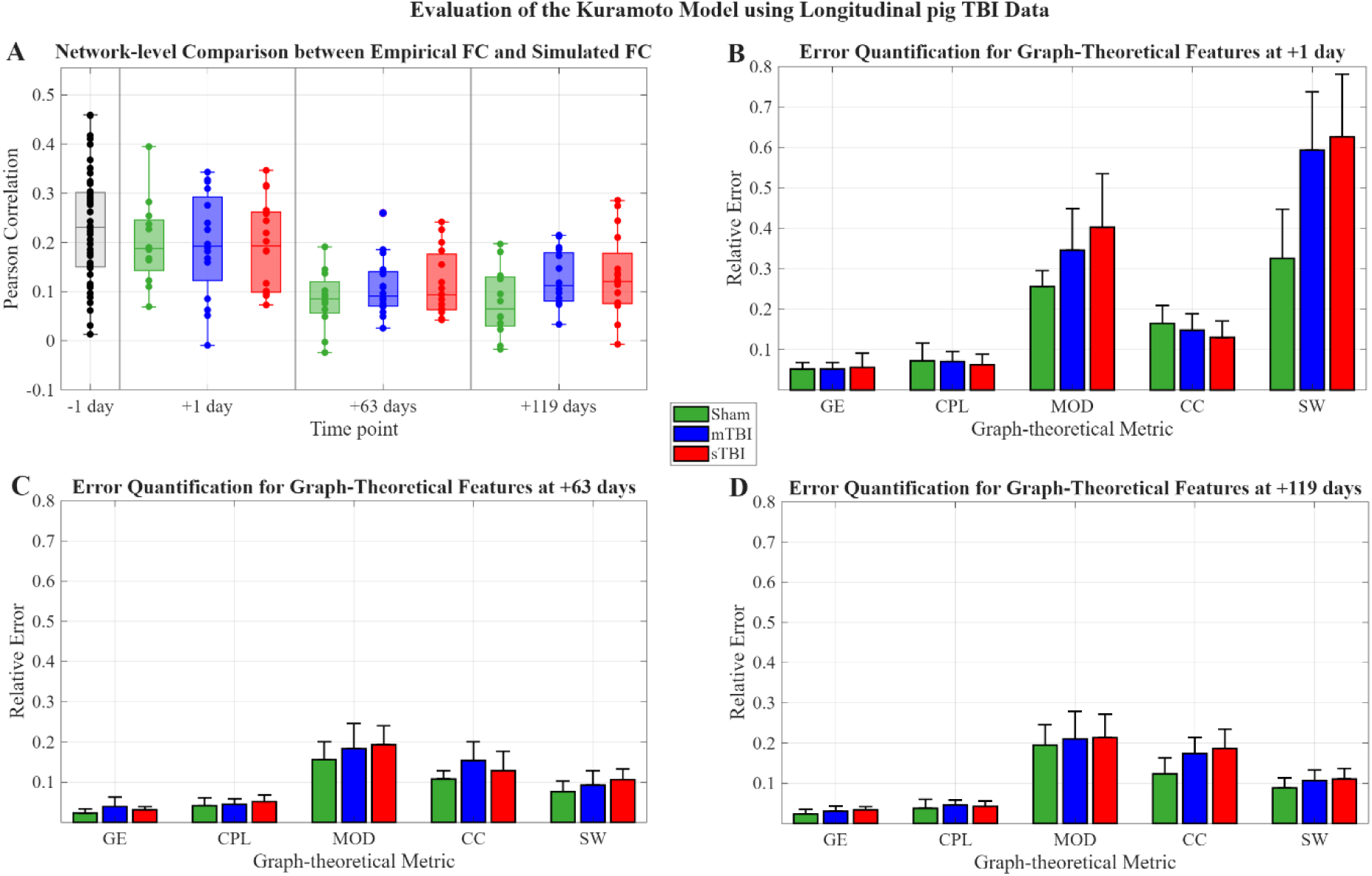
Performance evaluation of the Kuramoto model using longitudinal pig TBI dataset. (A) Distribution of network-level correlation between simulated and empirical FC across normal controls and TBI groups at +1 day, +63 days, and +119 days. Longitudinal error analysis for global efficiency (GE), characteristic path length (CPL), modularity (MOD), clustering coefficient (CC) and small worldness (SW) for sham, 3, and 9 mm TBI groups at +1 day, +63 days and +119 days are presented in 6B, 6C, and 6D respectively.

Graph theory metrics offer a comprehensive assessment of the model because they evaluated the underlying structural properties of the networks instead of just edge weights. Our findings showed a promising match between reproduced features in sFC and eFC using normal control pigs. We further evaluated the graph theory metric analysis to the longitudinal TBI dataset and summarized the assessment through relative error quantification. In **Figure 6**, relative errors for each metric presented for all TBI groups and time points. Immediately after the TBI (at +1 day, **Figure 6B**), quantified relative error in GE, CPL and CC were stable across the injury groups with averaged values of 0.05 for GE and CPL and ∼ 0.15 for CC. The quantified relative error in MOD and SW showed trends with injury severity; the least for the sham, moderate for the mild, and the highest for the severe TBI group. Furthermore, **Figure 6C** (at +63 days) showed an overall decline in the measured errors for all graph metrics with a substantial decline in SW and MOD. However, the trends identified for MOD and SW in **Figure 6B** remained the same in **Figure 6C** with a little difference. Relative error in other metrics including GE, CPL, and CC remained relatively stable. The graph theoretical features of eFC are observed to be reproduced in sFC at the +119 days (**Figure 6D**) and at the same extent as the +63 days. It is evident from the comparison of **Figure 6C** and **Figure 6D** that relative errors in all quantified metrics remained at the same level.

## Discussion

In the Kuramoto model framework, the natural frequencies and global coupling constant are determinant factors, instead of merely model parameters. The former is responsible for regional functional dynamics, and the latter represents structure influence on collective network dynamics (Kuramoto, 2005; Okuda & Kuramoto, 1991; Strogatz, 2000). Contrary to the findings of previous studies (Cabral et al., 2012; Lee et al., 2017; Wu et al., 2022), our data suggests that natural frequencies and global coupling constant are critically important for an accurate simulation of FC, and they should be carefully tuned using empirical data. In this context, we have tuned both parameters in a joint optimization algorithm using individual pig data and an averaged FC matrix of all normal control pigs. The quantified prediction score in **Figure 2A** (correlation between sFC and eFC) using individual data reached up to 0.347. The mean squared error also supported the findings via prediction scores such that it is smaller for higher prediction scores and vice versa. One of the important findings from this optimization, low prediction scores (high error) in the first row irrespective of *K* value (**Figure 2A** and **2B**) highlighted that randomly generated frequencies do not reflect natural frequencies of the brain oscillators. This is a direct validation that random distribution, especially those of less than 1Hz, is not a suitable choice for natural frequencies and identifying data-based optimal natural frequencies outperformed random distribution choice. Moreover, a global coupling constant of *K* > 10 seemed appropriate in controlling the influence of SC on brain oscillators to reproduce functional dynamics more accurately. In this situation, choosing an ideal *K* is difficult since a broad range of *K* show a stronger correlation and less variation. However, following the criterion in the methods section, an optimal value of *K* = 22 and individual natural frequencies for all pigs (distribution is presented in **Figure 2C**) were selected as individually tuned parameters. Alternatively, the heatmap in **Figure 2D** showed higher prediction scores (up to 0.55) and **Figure 2E** showed lower error values (with the least value of 0.1) with a tightly constrained range of *K* compared to the individual method. The prediction score and mean squared error in the averaged graph approach validated the irrelevance of randomly generated frequencies to the brain system and insufficient influence of SC at lower *K* values found in the individual method. In contrast to individual prediction scores, there are some irregularities (e.g., ups and downs of the difference measures against smaller *K* values) in **Figure 2D** since this tuning is based on an averaged matrix of all normal control pigs while **Figure 2A** is averaged after tuning through all pigs. Better prediction scores are the manifestation of cleaner and more stable FC data in contrast to individual FC graphs as shown in **Supplementary Figure S5**. In addition, functional graphs were highly heterogenous and SC graph were very consistent. Therefore, it is challenging for the Kuramoto model to reproduce clearly distinct FC patterns when input SC graphs were nearly identical. Considering this challenge, the importance of natural frequencies to reproduce individual FC patterns accurately becomes multifold. In **Figure 2F** tuned natural frequencies through averaged method are presented and separated into ipsilateral (red), contralateral (blue), and cerebellum (green) brain oscillators. We have identified a strong bilateral symmetry with similarity of 0.9201 between the ipsilateral and associated counterparts on contralateral hemisphere oscillators by quantifying correlation between the red and blue points in **Figure 2F**. The natural frequency values for oscillators were consistent with the distribution in **Figure 2C**. The set of tuned frequencies in **Figure 2F** may be helpful in capturing common FC patterns instead of individual pig specific FCs. In contrast, individually tuned frequencies target the pig-specific FC patterns in the population. In this study, we aimed to reproduce balanced FC patterns that may be achievable by using single global coupling constant, highly stable SC and pig-specific frequencies. It is important to note that individual tuning is computationally expensive and may not be an appropriate choice for some studies with very large datasets.

Moving forward, we next validated the chosen values of *K* for the individual and averaged approaches by examining the properties of Kuramoto order parameter; the synchrony and the metastability. In **Figure 3A**, the system of oscillators synchronizes as a function of *K* from 1-40 without reaching steady state. A possible stable synchronization was observed when 5 < *K* < 10, where synchrony was low and the model tended to approximate random patterns rather than meaningful network structure (Schmidt et al., 2015; Thümler et al., 2023). Additionally, metastability in **Figure 3B** suggested overall pig brains were more flexible to network state changes when 10 < *K* < 23. Based on the Kuramoto order parameter evaluation using the individual method, the pig brain seemed appropriately synchronized and with a flexible configuration at *K* = 22 that is consistent with prediction score findings. In comparison, synchrony curve in the averaged method conveys the impression of a steady state for *K* ≥ 20 with a synchrony value of ∼ 0.5. The metastability curve suggested an optimal *K* < 30 because the pig brain quickly tended towards rigid configuration beyond this point. An important finding from comparison of the individual and averaged approaches was that the average-based synchrony and metastability curves showed higher values, especially for 10 < *K* < 30. This suggested that the pig brain synchronizes better to capture common FC patterns accurately while system is yet flexible to accommodate changes in its network organization. Combining synchrony and metastability information, in averaged approach *K* = 29 was the optimal selection with high synchrony and metastability that also validates prediction score-based findings.

In the optimization process we have established that both averaged and individually tuned *K* and natural frequencies have specific advantages. Considering these advantages, a new combined approach (Combn) of global coupling constant from averaged tuning with individual tuned frequencies was developed. The Combn scheme (**Figure 4A**) showed a relatively better prediction score than the Indv and Avg schemes. Additionally, employing a single value of *K* was more intuitive to ensure a uniform global influence of SC on synchronization of oscillators in each pig brain network. Conversely, natural frequencies of oscillators may differ from one pig to another, indicating heterogeneity that can be preserved by using pig-specific natural frequencies. This variability among pig brains may potentially be suppressed by using a single set of tuned natural frequencies for each pig. In the light of these merits, we opted to use the common global coupling constant *K* = 29 from averaged tuning and pig-specific tuned natural frequencies to both reproduce common FC patterns and preserve heterogeneity. This answered one of our posed questions that combined use of common *K* and pig-specific tuned natural frequencies preserved the heterogeneity of the dataset. We used the combined parameters to simulate longitudinal functional networks of 3 post-TBI time points. We further validated our results by testing the observed correlation against a randomly generated null distribution. The empirical correlation was found to be significantly higher, demonstrating that the agreement between sFC and eFC was not driven by chance (**Figure 4B**). This result strengthens the evidence that the optimized model captures meaningful structure–function relationships rather than model-driven bias. Further analyses showed that the Kuramoto model accurately reproduced FC patterns, which were evident from a strong direct relationship (*r* = 0.6081, *p* < 0.001, **Figure 4C**) and visual inspection between the edges of averaged eFC (**Figure 4D**) and sFC (**Figure 4E**).

In terms of graph-theoretical properties, **Figure 5 (A–E)** demonstrated that the Kuramoto-based simulations capture certain topological features more faithfully than others. The strongest correspondence for GE (**Figure 5A**) suggested that the large-scale integrative capacity is primarily constrained by the underlying structural architecture. This was consistent with prior findings, indicating that efficiency-related measures are closely tied to anatomical path structure and connection density (Honey et al., 2007; Rubinov & Sporns, 2010). Next, CPL and CC (**Figure 5B and 5D**) also exhibited comparable trends between empirical and simulated network features with small deviations, indicating that local segregation and integration are substantially, but not exclusively, shaped by structural connectivity. In contrast, MOD and SW (**Figure 5C and 5E**) displayed larger discrepancies between the empirical and simulated networks. These properties reflected higher-order organizational principles that depend not only on static anatomical wiring, but also on dynamic coordination patterns and state-dependent processes (Deco et al., 2011; Sporns, 2013). Further, the error analysis (**Figure 5F**) supported this interpretation, suggesting that integrative metrics were more directly shaped by SC, whereas higher-order organization may require additional dynamical or contextual factors beyond static connectivity alone.

Extending the network-based analysis to the longitudinal experimental TBI dataset allowed us to examine how injury may influence structure–function interaction (**Figure 6A**). Immediately after injury (at +1 day), correlation distributions were comparable to those of controls (−1 day), irrespective of injury severity. It suggested that acute structural perturbations may not immediately disrupt large-scale coordination mechanisms, or that functional compensatory processes immediately preserve functional organization despite focal damage (Hillary & Grafman, 2017; Sharp et al., 2014). In contrast, we observed a progressive reduction in network-based correlation at the +63 days and +119 days across all three groups. This decline reflected the potential gradual network reorganization and alterations in intrinsic oscillatory properties (i.e., natural frequency) over time. These potential factors were found to be consistent to previously published studies on TBI, reported evolving changes in white matter integrity and excitatory–inhibitory balance, which modified regional intrinsic synchronization dynamics (Johnson et al., 2013; Sharp et al., 2014). Additionally, some studies on humans reported alterations in natural frequencies of brain regions (Gómez et al., 2013; Marshall et al., 2002). This shift in natural frequencies has weakened the structure–function coupling captured by the Kuramoto model. Similar findings of reduced structure-function correspondence were reported by recently published studies (Liu et al., 2024; Y. Sun et al., 2024). Importantly, no significant differences were detected across groups at corresponding time points, suggested that our model has maintained prediction performance across injury severities. Combining cross sectional and longitudinal findings, indicated that while TBI-related adaptations and temporal effects may modulate intrinsic dynamics, the large-scale dynamical principles linking structural architecture to functional organization remain relatively robust.

Moreover, at +1 day (**Figure 6B**), relative error values were low for GE and CPL across all groups. This indicated that integrative network properties remain closely constrained by the underlying structural topology in the acute phase of TBI. The errors in MOD and SW were comparatively larger, especially increasing with TBI severity. This pattern was found to be consistent with prior work showed modular organization and small-world properties were more sensitive to dynamic coordination and pathological perturbations (Hillary & Grafman, 2017; Sharp et al., 2014). At +63 days (**Figure 6C**) and +119 days (**Figure 6D**), relative errors for GE and CPL remain consistently low across all groups, validated integrative properties were structure-driven even over time. Errors in MOD and CC show reduced separation between groups compared to the +1 day, and SW demonstrates relatively small longitudinal variation. This temporal stabilization reflected adaptive reorganization or plasticity mechanisms following TBI (May, 2011; Zilles & Amunts, 2015). Overall, the persistence of low error in integration-based metrics across all time points supported the notion that large-scale structural architecture continues to govern core functional organization, while higher-order topological features exhibited greater sensitivity to TBI-related and time-dependent network adaptations (Deco et al., 2011).

## Limitations

This study has several limitations. First, although the swine brain provides a valuable approximation of human neuroanatomy, species differences may still limit the direct translation of findings from swine TBI to human TBI. Second, the sample size was modest, which may reduce statistical power, especially for detecting subtle or long-term effects. Third, the injury model employed here produced controlled focal impacts, which may not fully capture the heterogeneity of TBI mechanisms seen clinically. Finally, the follow-up duration was only 4 months, limiting assessment of chronic pathological changes. Future studies should incorporate larger sample sizes, expanded injury paradigms, and longer observation periods to strengthen and extend the present findings.

## Conclusion

This study demonstrated that a structurally constrained and jointly optimized Kuramoto model can reliably reproduce key functional connectivity patterns in the pig brain network. Careful calibration of natural frequencies and global coupling constant was essential for achieving meaningful structure–function correspondence, thus emphasizing the mechanistic interplay between regional dynamics and anatomical wiring. The model robustly captured integration-related graph metrics such as global efficiency, characteristic path length, and clustering, while more complex organizational features showed greater sensitivity to TBI and temporal effects. Importantly, the prediction performance remained comparable across mTBI and sTBI groups and following longitudinal stages, suggesting that fundamental structure-driven coordination mechanisms persist despite injury.

From a translational perspective, this framework provides a quantitative tool to probe how an TBI reshapes functional dynamics and may help identify mechanistically grounded biomarkers of following recovery or maladaptation. The future work will extend longitudinal optimization of model parameters to track injury-specific dynamical changes and evaluate their potential utility in guiding therapeutic interventions.

## Acknowledgements

This study was funded by the National Institutes of Health (NIH) grants R21NS123732 and R21NS131526.

## Declaration of Generative AI tools in the writing process

During the preparation of this work the author(s) used AI tools (i.e., ChatGPT, NotebookLM, Copilot) in order to improve language. After using this tool/service, the author(s) reviewed and edited the content as needed and take(s) full responsibility for the content of the publication.

## Author Contributions

Ishfaque Ahmed: Writing – review & editing, Writing – original draft, Visualization, Software, Methodology, Investigation, Formal analysis. Morgan H. Laballe: Software, Methodology. Moira F. Taber: Writing – review & editing, Data curation. Sydney E. Sneed: Data curation. Erin E. Kaiser: Writing – review & editing, Resources, Project administration, Funding acquisition, Data curation, Conceptualization. Franklin D. West: Writing – review & editing, Resources, Project administration, Funding acquisition, Conceptualization. Taotao Wu: Writing – review & editing, Resources, Methodology. Qun Zhao: Writing – review & editing, Validation, Supervision, Resources, Methodology, Funding acquisition, Conceptualization.

