## Supplemental Material for "Assessment of Coupled Phase Oscillators-Based Modeling in Swine Brain Connectome"

*Supplementary TABLE 1. Focused brain regions in this study and parcellated from previously published atlas by Saikali et al., (2010).*

| Brain region | Abbreviation | Right | Left | Cerebellum |
|---|---|---|---|---|
| Ventral Anterior Thalamic Nucleus | VATN | 1 | 27 | - |
| Caudate Nucleus | CN | 2 | 28 | - |
| Putamen | PUT | 3 | 29 | - |
| Hippocampus | HIP | 4 | 30 | - |
| Amygdala | Amy | 5 | 31 | - |
| Primary Somatosensory Cortex | PSC | 6 | 32 | - |
| Primary Motor Cortex | PMC | 7 | 33 | - |
| Somatosensory Association Cortex | PAC | 8 | 34 | - |
| Premotor Cortex | PC | 9 | 35 | - |
| Dorsolateral Prefrontal Cortex | DPC | 10 | 36 | - |
| Anterior Prefrontal Cortex | APC | 11 | 37 | - |
| Insular Cortex | IC | 12 | 38 | - |
| Primary Visual Cortex | PVC | 13 | 39 | - |
| Secondary Visual Cortex | SVC | 14 | 40 | - |
| Associative Visual Cortex | AVC | 15 | 41 | - |
| Inferior Temporal Gyrus | ITG | 16 | 42 | - |
| Middle Temporal Gyrus | MTG | 17 | 43 | - |
| Superior Temporal Gyrus | STG | 18 | 44 | - |
| Dorsal Posterior Cingular Cortex | DPCC | 19 | 45 | - |
| Dorsal Anterior Cingulate Cortex | DACC | 20 | 46 | - |
| Anterior Entorhinal Cortex | AEC | 21 | 47 | - |
| Peri Rhinal Cortex | PRC | 22 | 48 | - |
| Para Hippocampal Cortex | PHC | 23 | 49 | - |
| Fusiform Gyrus | FG | 24 | 50 | - |
| Auditory Cortex | AC | 25 | 51 | - |
| Pre-Pyriform Area | PPC | 26 | 52 | |
| Cerebellar Lobule III | CL3 | - | - | 53 |
| Cerebellar Lobule IV | CL4 | - | - | 54 |
| Cerebellar Lobule V | CL5 | - | - | 55 |
| Cerebellar Lobule VI | CL6 | - | - | 56 |
| Cerebellar Lobule VII | CL7 | - | - | 57 |
| Cerebellar Lobule VIIIB | CL8B | - | - | 58 |
| Crus - I | CR1 | - | - | 59 |
| Crus - II | CR2 | - | - | 60 |

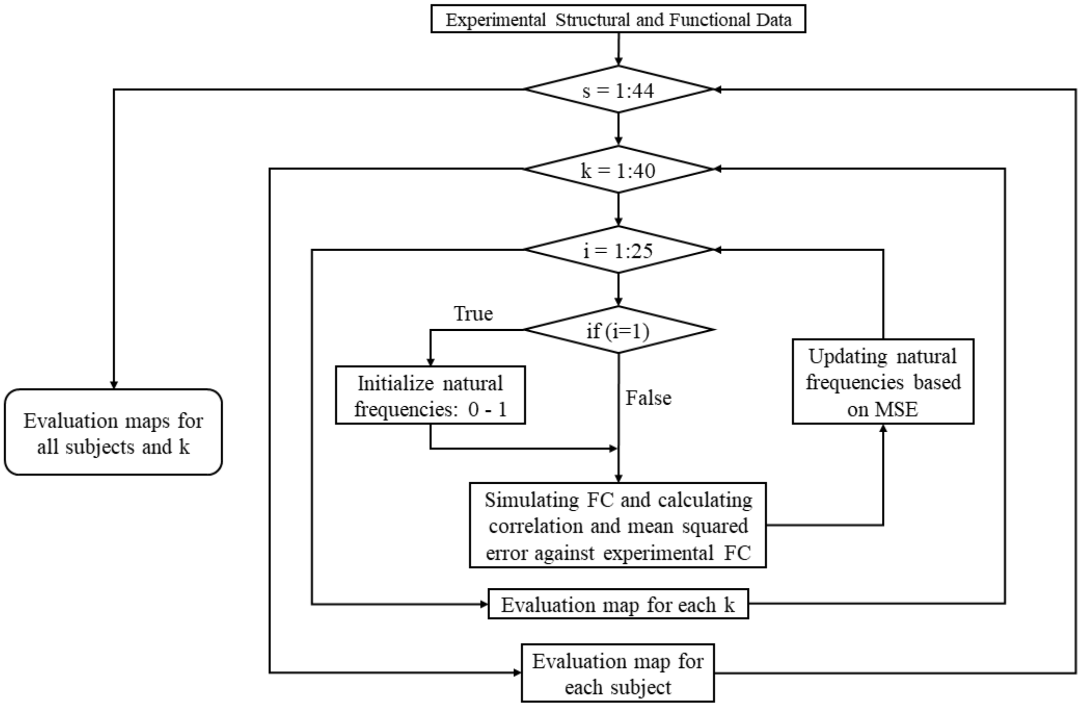


*Supplementary Figure S1. Representation of the parameter optimization framework. There are 44 normal subjects that are used to optimize the best coupling constant and natural frequencies for the nodes/oscillators. To fully evaluate the models' capabilities, we have tested the parameters on each individual and the average of 44 normal subject data.*

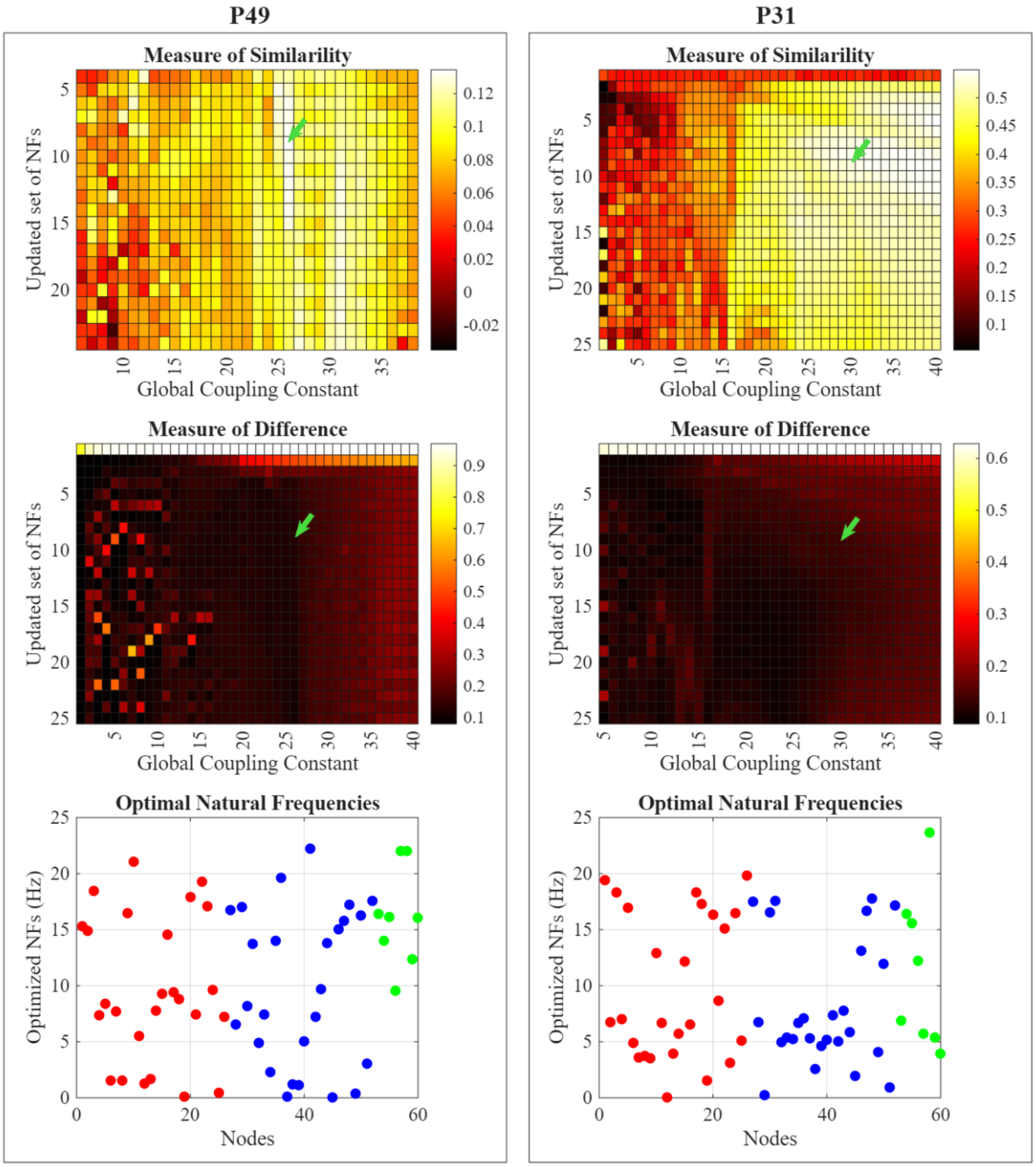


*Supplementary Figure S2. Showing individual tuning similarity and difference maps and tuned natural frequencies. These similarity and difference maps are constituent maps of the results shown in Figure 2(A–B). In this approach, global coupling constant and natural frequencies were selected using the same procedure for each individual.*

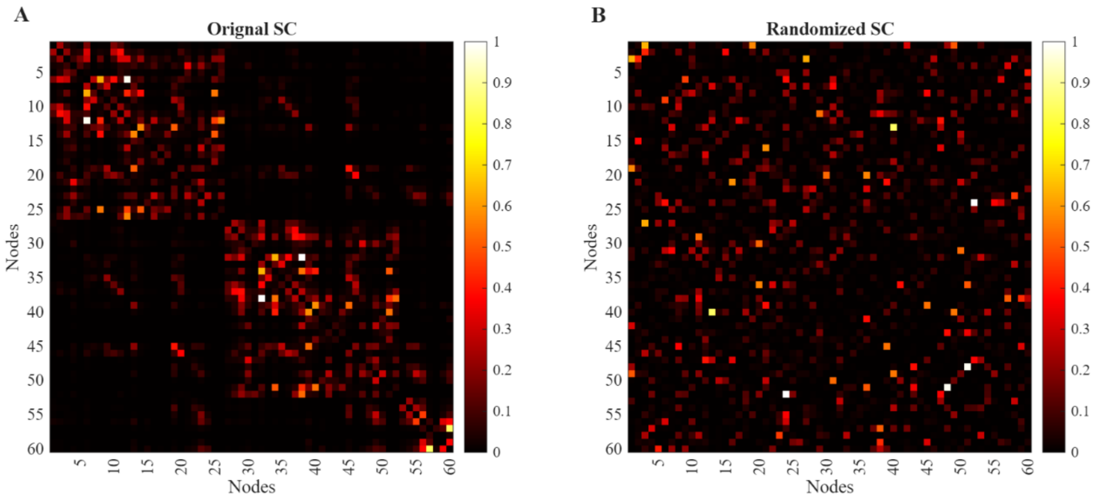


*Supplementary Figure S3. Heatmaps showing original structural connectivity matrix for a subject (A) and its shuffled/randomized copy (B).*

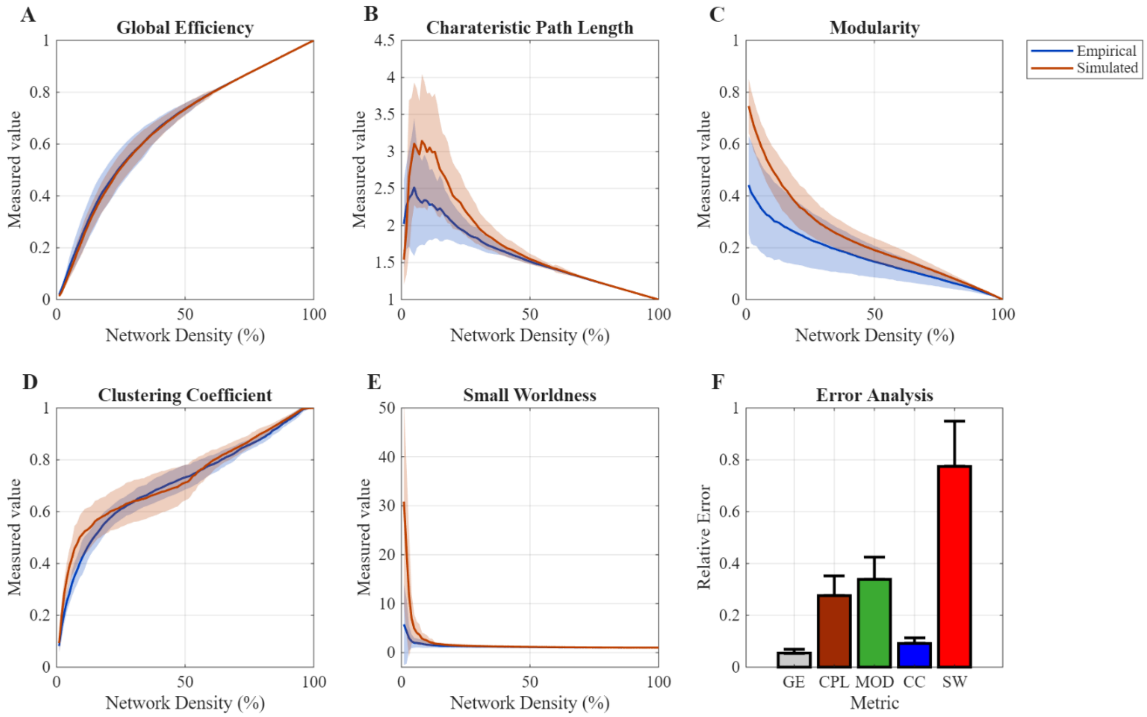


*Supplementary Figure S4. Evaluation of the Kuramoto model based on graph theoretical features in the range of 1% to 100% network densities. Overlay of the mean and distributions of empirical and simulated global efficiency (A), characteristic path length (B), modularity (C), clustering coefficient (D) and small worldness (E). F shows the measure of relative error for all the five metrics.*

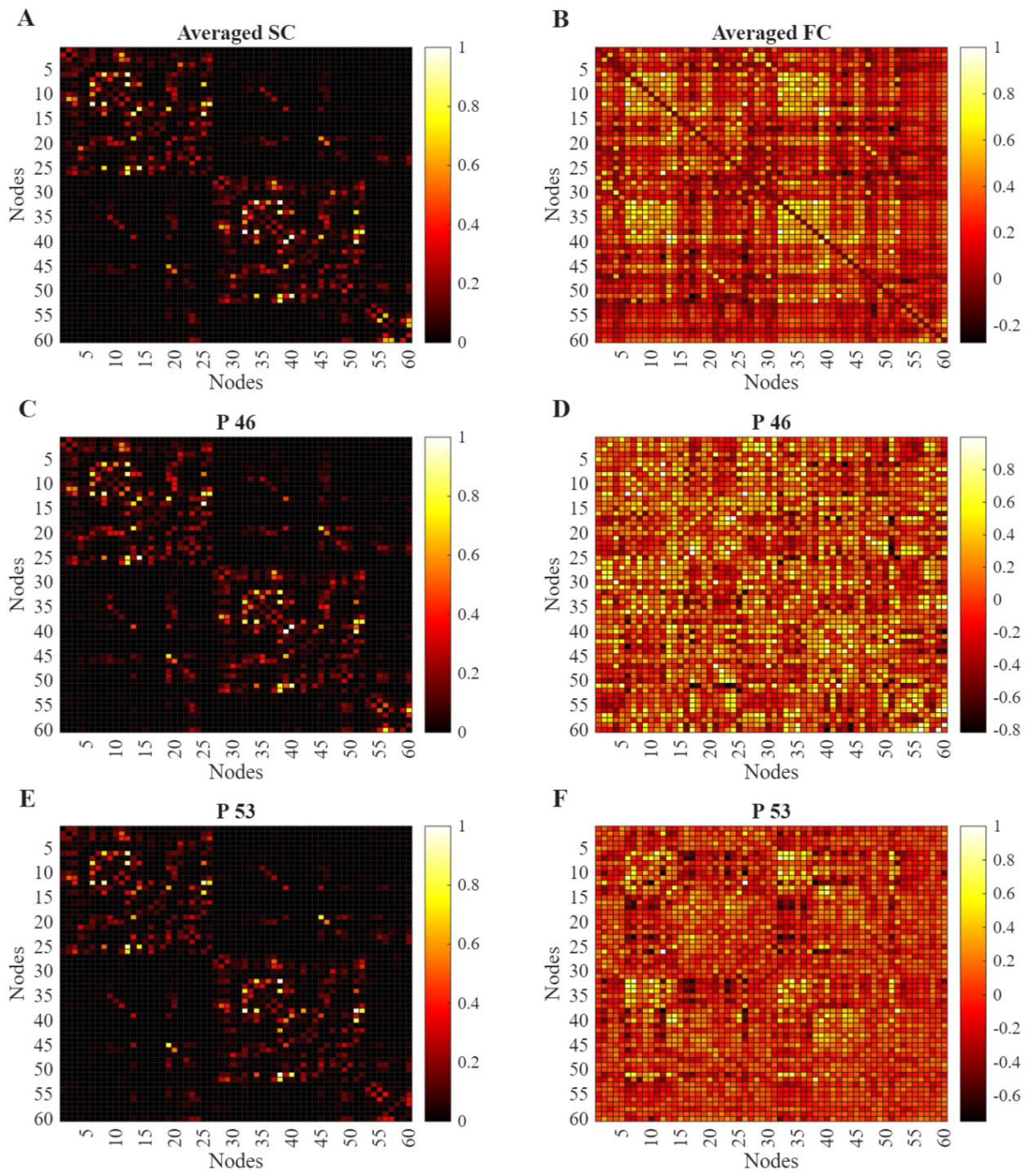


*Supplementary Figure S5. Illustration of inter-subject variability identified across pig brain networks.* ***Left Column****: A shows the averaged structural connectivity (SC) across 44 pigs, C and E show SC networks of two pigs.* ***Right Column****: B shows the averaged functional connectivity (FC) across 44 pigs, D and F show SC networks of two pigs.*

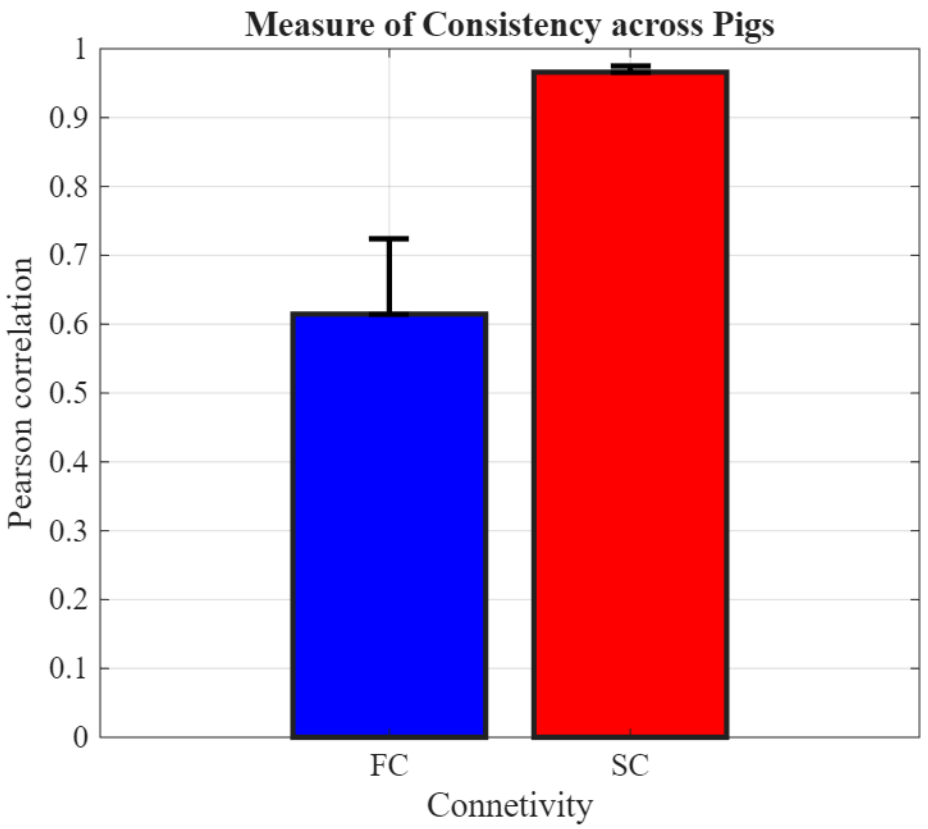


*Supplementary Figure S6. Quantified difference of functional connectivity (FC) and structural connectivity (SC) consistency across normal control subject.*

### Appendix A

**Global efficiency**: It is a graph theory feature that represents how efficient information is processed between nodes. It is measured as the average of inverse of shortest path lengths.

**Characteristic path length**: The average shortest path length represents the characteristic path length. The higher values indicate slow information processing and vice versa.

**Modularity**: It measures the degree to which the network can be divided into communities with dense intra-module connections and sparse inter-module connections.

**Global clustering coefficient**: The clustering coefficient quantifies the tendency of nodes in a network to form locally interconnected groups. It measures the extent to which the neighbors of a node are also connected to each other. It is reflecting the level of local segregation and redundancy within the network.

**Small worldness**: Small worldness quantifies the balance between network segregation and integration relative to a comparable random network. A network is said to be a small world by high clustering among neighboring nodes while maintaining short average path lengths which allows efficient information transfer across the system.
